# Sexual dimorphism in pollen foraging and sensory traits in *Heliconius* butterflies

**DOI:** 10.1101/2025.09.18.676771

**Authors:** José Borrero, David F. Rivas-Sánchez, Daniel Shane Wright, Caroline Nicole Bacquet, Stephen H. Montgomery, Alexander Keller, Richard M. Merrill

**Affiliations:** Division of Evolutionary Biology, Faculty of Biology, LMU Munich, Germany; School of Biological Science, University of Bristol, Bristol, United Kingdom; Universidad Regional Amazónica de Ikiam, Tena, Ecuador; Cellular and Organismic Networks, Faculty of Biology, LMU Munich, Germany

**Author notes:** Corresponding authors (J.B.) & (R.M.M).

## Abstract

Sexual dimorphism in foraging behaviour is widespread in insects and may arise from differences in nutritional demands, sensory systems, or cognition. *Heliconius* pollen-feeding is an evolutionary innovation among butterflies that supports extended lifespans and sustained reproduction. However, how foraging behaviour varies between the sexes and how it relates to sexually dimorphic traits remains poorly understood. We investigated sex-specific foraging strategies in wild *Heliconius himera*, a highland specialist from southern Ecuador, using field surveys and DNA metabarcoding. Females carried more pollen than males, consistent with higher nutritional demands, yet we observed no differences in plant richness and composition. This indicates that sex differences reflect effort in foraging behaviour rather than shifts in plant choice. Gut samples revealed greater pollen diversity and a more consistent community profile than proboscis samples, suggesting they better capture cumulative foraging history. We also quantified sexually dimorphic sensory traits and found that males had larger eyes and more ommatidia, whereas females had larger mushroom bodies. While the functional significance of these differences remains unclear, these patterns are consistent with sexual dimorphism reported across *Heliconius* and suggest males and females may be under divergent selective pressures. Our findings highlight how sex-specific foraging differences can arise from differential effort on shared floral resources and co-occur with divergent sensory and neural investment, offering insights into the ecological basis of intraspecific variation in pollen use.

## Introduction

Sexual dimorphism is widespread across animals and often involves divergence in morphology, physiology, and behaviour. Among insects, sex differences in foraging behaviour may arise from distinct dietary needs or sensory biases. Many species exhibit sex-specific specializations in visual and olfactory systems that support different ecological roles, such as mate location in males and resource acquisition in females (Smith et al. 2019). For instance, in the oil bee *Macropis fulvipes*, both sexes forage on the same flowers, but males rely more on visual cues and only collect pollen, while females show a stronger preference for olfactory cues and collect both pollen and floral oils (Dötterl et al. 2011). Similarly, in the fly *Bibio rufiventris*, males and females differ in their responses to floral volatiles over the course of the day, reflecting sex-specific attraction to scent cues (Zhu et al. 2025). Despite such examples, the ecological and sensory drivers of sex-specific foraging behaviour remain poorly understood in most insect taxa.

Pollen-feeding is a widespread and ecologically important behaviour in insects, particularly among bees, flies, and beetles, where it plays a central role in adult nutrition, reproduction, and pollination (Romeis et al. 2005). These lineages have evolved diverse morphological and behavioural adaptations to collect and digest nutrient-rich pollen (Roulston and Cane 2000; Ruedenauer et al. 2019). Unlike most butterflies that feed only on nectar, *Heliconius* uniquely show a derived pollen-feeding behaviour, a strategy central to their nutrition, reproduction, and mutualism with the plants they pollinate (Gilbert 1972; Young and Montgomery 2020). Sex differences in pollen feeding have been documented in some *Heliconius* species (Boggs et al. 1981; Cardoso 2001; Mendoza-Cuenca and Macías-Ordóñez 2005), but how these differences align with sexually dimorphic sensory traits, and whether they involve shifts in pollen load versus composition, remains understudied.

Insects that feed on pollen gain access to amino acids and lipids otherwise scarce in nectar (Romeis et al. 2005). In *Heliconius*, pollen feeding has been linked to enlarged mushroom bodies that support sensory integration and learning, possibly facilitating this unique behaviour (Sivinski 1989; Montgomery et al. 2016). Both sexes benefit from pollen-derived nutrients that extend lifespan and sustain reproduction (Dunlap-Pianka et al. 1977; Pinheiro de Castro et al. 2020), but the direct benefits differ: in females, amino acids and lipids directly support oogenesis and sustained egg production, and in older females, pollen-feeding helps maintain adult cyanogenic defences that contribute to Müllerian mimicry (O’Brien et al. 2003; O’Brien et al. 2004; Pinheiro de Castro et al. 2025). In males direct benefits appear more limited, where it mainly supports spermatophore production and mating effort (Boggs and Gilbert 1979; Boggs et al. 1981). These differences in direct benefits likely drive a female bias in pollen foraging.

Multiple studies have found evidence for sex differences in pollen feeding in *Heliconius*. Field observations of *H. charithonia*, a species within the *erato/sara/sapho* clade, have revealed sex-specific pollen use: females are more often recorded on *Hamelia patens* plants, while males favor *Lantana camara* (Mendoza-Cuenca and Macías-Ordóñez 2005). Similarly, broader studies across *Heliconius* have documented larger pollen loads and longer foraging duration in females than males (Boggs et al. 1981; Gilbert 1991; Cardoso 2001). These patterns mirror broader trends in other Lepidoptera, where females actively seek out nectar with higher amino acid content than males (Alm et al. 1990; Erhardt and Rusterholz 1998; Mevi[Schütz and Erhardt 2005).

In *Heliconius*, differences in foraging may be influenced by sexually dimorphic visual systems. Unlike most butterflies, *Heliconius* possess two UV-sensitive opsins, and in the erato/sara/sapho clade one of these (UVRh1) is linked to the sex chromosome (W) and expressed only in females, enabling UV wavelength discrimination in females, which may enhance floral detection capabilities (Finkbeiner and Briscoe 2021; Chakraborty et al. 2023). However, the ecological relevance remains unclear; for instance, UV sensitivity does not appear to affect oviposition in *H. erato* and *H. himera* (Borrero et al. 2023). Beyond opsin expression, other visual traits such as facet number, facet diameter, and corneal area shape acuity and sensitivity and can differ between sexes (Land 1997; Wright et al. 2023). As such, integrated studies of foraging behaviours and sensory traits may help elucidate the basis of sex-specific pollen foraging.

*Heliconius himera* is a highland specialist inhabiting seasonally dry forests of southern Ecuador (∼1500 m asl). Compared with its lowland relatives, it occupies more open habitats with higher light intensity, stronger temperature fluctuations, and distinct floral communities, conditions that likely shape both resource availability and foraging strategies (Descimon and Mast 1984; Jiggins et al. 1996; Dell’Aglio, Mena, et al. 2022). Reflecting adaptation to this environment, *H. himera* differs from lowland *H. erato* in brain morphology and sensory processing, investing more in antennal lobes and less in visual neuropils (Montgomery and Merrill 2017). When visual and olfactory cues conflict, *H. himera* relies more heavily on olfactory input than *H. erato* (Dell’Aglio, McMillan, et al. 2022), a pattern also seen in other Lepidoptera with variable neural investment (Stöckl et al. 2016; Borrero et al. 2024). As an incipient species within the *erato/sara/sapho* clade, *H. himera* likely shares the sexually dimorphic opsin expression and vision reported in related taxa (McCulloch et al. 2017; Chakraborty et al. 2023). This combination of distinct sensory weighting and potential sexually dimorphic vision makes *H. himera* a compelling model for exploring how environmental adaptation and sex-specific sensory traits shape foraging behaviour.

Despite its ecological importance, pollen use in *Heliconius* remains incompletely understood. Traditional methods such as field observations and palynological keys have revealed diverse diets dominated by *Psychotria*, *Psiguria*, and *Lantana* (Boggs et al. 1981; Cardoso 2001; Estrada and Jiggins 2002; Mendoza-Cuenca and Macías-Ordóñez 2005). However, these methods are slow, expertise-dependent, and limited in taxonomic resolution. DNA metabarcoding provides a powerful alternative: targeting regions such as ITS2, enables high-throughput, species-level identification of mixed pollen samples (Keller et al. 2015; Bell et al. 2016). Already widely applied in bee pollination studies (Piko et al. 2021; Wilson et al. 2021; Martins et al. 2023), this method can open new perspectives on *Heliconius* foraging ecology and sex-specific pollen use.

In this study, we integrate field surveys, DNA metabarcoding, and morphological analyses to investigate sex differences in foraging behaviour in *H*. *himera*. Specifically, we tested the hypothesis that sexes differ in the amount and diversity of pollen collected, reflecting distinct nutritional demands or foraging effort. We also quantified eye and brain morphology to document sexually dimorphic sensory and neural traits. This study provides one of the first molecular characterizations of pollen resource use in *Heliconius* and highlights how sex-specific differences in pollen collection can co-occur with divergent sensory and neural investment.

## Materials & Methods

### Study sites

Field surveys were conducted at five sites in Loja Province, Ecuador, around the city of Vilcabamba (4°15′S, 79°13′W), at elevations of 1500–1700 m (Fig. 1A–B). All sites are characterized by thorn scrub dry forest dominated by *Acacia macracantha* and xerophytic vegetation (Jiggins et al., 1996). Rumi Wilco is a 40-ha private reserve containing ∼700 plant species (Jorgensen & Leon-Yanez, 1999), while Cerro Mandango is a nearby mountain site dominated by Acacia. The other sites (Quebrada Puliche, Agua de Hierro, and San Pedro) occur along small streams with similar vegetation, adjacent to coffee plantations and pastures.

**Figure 1:**
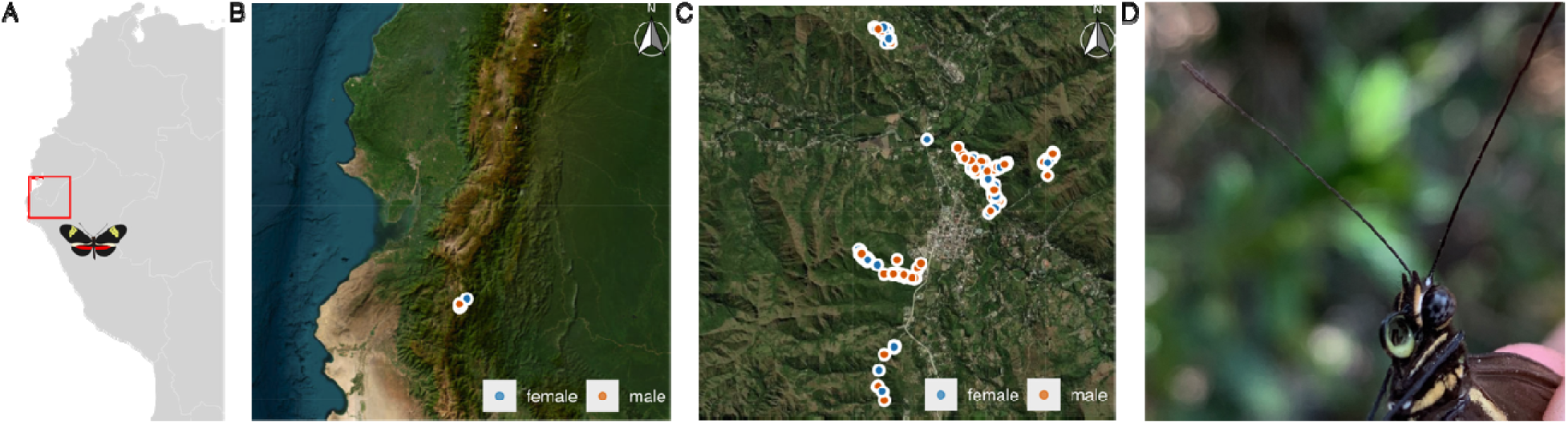
Sampling region and pollen foraging behavior in H. himera. (**A**) Study area in southern Ecuador. (**B**) Overview map showing the location of butterfly survey sites in Loja Province. Points represent individual collection records, with colour indicating sex (blue = female, orange = male). (**C**) Satellite map of survey sites, including Rumi Wilco, Cerro Mandango, Quebrada Puliche, and San Pedro. (**D**) Close-up photograph of a H. himera with a visibly pollen-loaded proboscis.

### Butterfly sampling

From April to September 2023 spanning the wet-to-dry season transition, we sampled *H. himera* along trails at all five sites, with most effort concentrated at Rumi Wilco and Cerro Mandango, which were visited at least five days per week. Daily sampling took place between 08:00 and 17:00. This sampling window corresponds to the main period of *Heliconius* activity: *H. himera* is active early in the morning and remains active into the late afternoon (Davison et al. 1999) and *Heliconius* species in Panama also show sustained activity across 08:00–16:00 (Dell’Aglio et al. 2024). Recording the exact capture time, allowed us to examine potential sex differences in activity pattern.

Each butterfly was captured, marked on the forewing, and released after recording GPS location, time of capture, sex, body length, pollen load and observer identity. Pollen load was visually classified into categories based on the proportion of the proboscis covered with pollen: none (0%), small (<50%), and large (>50%) (Fig. 1D). Two observers collected these data, and observer identity was recorded for all individuals to account for potential observer effects. Our categorical scoring system is comparable to pollen-load ranking approaches used in previous *Heliconius* field studies (Boggs et al. 1981; Cardoso 2001; Mendoza-Cuenca and Macías-Ordóñez 2005). Recaptured individuals were noted for movement analyses. In total, 877 butterflies were recorded (371 females, 506 males).

Microclimate measurements were collected for a subset of butterflies during routine fieldwork. Temperature (°C) and relative humidity (%) were recorded at capture sites using HOBO Temp/RH U23-002 data loggers placed opportunistically on days when loggers were available. Loggers were deployed directly at each capture location and recorded continuously for at least 24 h post-capture. The 49 individuals with microclimate measurements (24 females, 25 males) represent an opportunistic subset determined by logger availability, collected between 26 April and 12 September 2023, spanning 20 sampling days across April, May, June, July, August, and September. Sex was sampled randomly across this period, with males and females collected throughout the full sampling period. These individuals did not overlap with the butterflies used for pollen metabarcoding, and microclimate data were used only for exploratory analyses of microhabitat differences between sexes.

### Pollen collection and ITS2 sequencing

To identify pollen sources, we collected 71 *H. himera* (25 females, 46 males) with hand nets between July and August 2023. The sample contained more males than females because, as is common in *Heliconius* field surveys, males are more frequently encountered and captured (Boggs et al. 1981; Cardoso 2001; Jiggins 2017). Additionally, a subset of females was retained alive to maintain stock cages for parallel experiments. Specimens were placed in entomological envelopes; proboscises were dissected and stored in PCR tubes, while bodies (containing guts) were preserved in DMSO/EDTA/NaCl solution. Samples were kept at −15 °C until DNA extraction and sequencing in February 2024. All proboscises (25 females and 46 males), and a subset of guts (13 females, 26 males) were analysed using ITS2 DNA metabarcoding, each as an individual sample (not pooled). DNA metabarcoding enables the identification of pollen to the genus or species level from mixed samples, without the need for specialized expertise in traditional palynology (Keller et al. 2015).

DNA was extracted using the Macherey–Nagel NucleoSpin Food Kit (Düren, Germany), exactly following the ITS2 metabarcoding protocol of Sickel *et al*., (2015), with additional practical details on reagents and laboratory steps drawn from (Campos et al. 2021) (Section 8: Standard method for identification of bee pollen mixtures through meta-barcoding). We included nine negative controls (DNA-free molecular-grade water) and four positive controls (pre-established 1:3 mixtures of *Musa acuminata* and *Mangifera indica* plant DNA) throughout laboratory processing to detect potential contamination. PCR amplification, dual-indexing, library preparation, and sequencing were performed in-house at LMU. Sequencing was carried out on an Illumina MiSeq using the MiSeq Reagent Kit v2 (2 × 250 bp) (Illumina Inc., San Diego, CA, USA), with custom read and index primers added to the reagent cartridge following Campos et al. 2021.

Raw sequence reads were processed combining open-source tools with a custom bioinformatics pipeline (https://github.com/chiras/metabarcoding_pipeline). Reads were demultiplexed, primer-trimmed, and merged using VSEARCH (Rognes et al. 2016), with short reads (<150 bp) discarded. Amplicon sequence variants (ASVs) were inferred via dereplication and denoising, and chimeric sequences removed. Taxonomic assignments were performed via global alignment (at least 97% identity threshold, best hit selection) to a curated regional ITS2 plant reference database built using BCdatabaser (Keller et al. 2020). Unclassified reads were further annotated first with global alignments against a curated worldwide database (Quaresma et al., 2024), then if still unclassified with the same database hierarchically using the SINTAX algorithm as taxonomically deep as possible (Edgar 2016). Taxa were retained if they reached ≥1% relative abundance. We restricted analyses to vascular plants using a curated Streptophyta reference database (Quaresma et al. 2024) that excludes algae and fungi.

Sequencing generated a mean raw throughput of 30,401 ± 14,445 reads per sample (range: 170–101,135, including negative controls). After quality filtering, merging, dereplication, chimera removal, and taxonomic assignment, samples retained an average of 12,175 reads per sample (range: 141–31,662). Negative controls showed only very low read counts, while positive ITS2 mock controls amplified as expected, confirming run quality and supporting the removal of potential contaminants. All remaining samples retained sufficient read depth for downstream ecological analyses.

### Wing reflectance spectrometry and age estimation

To estimate the relative age of wild individuals, we adapted a spectrometric method developed by Dell’Aglio et al. (2017), who used wing colour metrics to infer butterfly age. We collected full-spectrum reflectance data collected from the red and yellow wing bands of insectary-reared individuals of known age. Reflectance spectra were measured using a Flame-T-XR1-ES Miniature Spectrometer (Ocean Optics Inc.) connected to a PX-2 Pulsed Xenon light source (spectral range 220-750 nm) and an Ocean Optics OCF-109053 UV/Vis bifurcated reflection probe with 200µm fibre diameter. All reflectance measurements were standardized with a WS-1 diffuse white reflectance standard. For each recording (integration time: 2500 ms; three-scan average), the probe was positioned at a 45° angle and held 1 mm from the wing surface using an in-house probe mount. This configuration produced an illuminated sampling spot approximately 2 mm in diameter. For each individual, we collected three replicate spectra per colour patch (red or yellow), each taken from a different spot within the patch to capture within-patch variation.

Reflectance was analysed in R with the pavo package (Maia et al. 2019), and the area under the reflectance curve (AUC) was calculated for each colour patch. In total, we measured reflectance and age data for 320 insectary-reared individuals across seven *Heliconius* species with red and/or yellow coloration, including 108 insectary reared *H. himera.* Age was calculated from known eclosion and sampling dates and linked to reflectance data via standardized individual identifiers.

Age was predicted using linear models with log-transformed AUC values as predictors. A combined model including both red and yellow AUCs provided the best fit (adjusted R² = 0.13, P < 0.001) and was used to estimate the age of wild *H. himera*. Predicted values were back-transformed to age in days, and each individual was assigned a relative age score. This continuous relative age metric provided a standardized measure across individuals and was used to test how pollen foraging behaviour varies with age in the wild population.

### Eye cuticle morphology

To test for sexual dimorphism in eye cuticle morphology, we analysed wild-caught and insectary-reared *H. himera* following established protocols (Wright et al. 2024; Wright and Merrill 2025a; Wright and Merrill 2025b). Eyes were dissected and incubated in 20% sodium hydroxide (NaOH) for 18–24 hours to loosen internal tissues beneath the corneal cuticle. After incubation, each eye was carefully cleaned, mounted in Euparal (Carl Roth GmbH) on a microscope slide, and left to cure overnight.

Slides were imaged at 25× magnification using a Leica M80 stereomicroscope equipped with a Flexacam C1 camera and Leica Application Suite X software. All images included a 1 mm scale bar for size calibration. Corneal surface area was measured using the Freehand Selection and Measure tools in ImageJ/Fiji (Schindelin et al. 2012), while total ommatidia (facet) number was estimated via image thresholding and the Analyze Particles function. To account for variation in body size, we additionally measured inter-ocular distance and hind tibia length using the Straight-Line tool. Based on the strong bilateral symmetry previously reported for these traits (Wright et al. 2024), we measured the left eye for all analyses unless it was damaged or missing, in which case the right eye was used.

### Brain immunohistochemistry and Imaging

We examined sexual dimorphism in brain morphology in 43 *H. himera* (21 females, 22 males), combining previously analysed wild individuals from Vilcabamba (8 females, 8 males) and two cohorts of common-garden reared butterflies, from Cambridge (5 females, 5 males)(Montgomery and Merrill 2017) and Tena, Ecuador (8 females, 9 males). All insectary-reared butterflies (both Cambridge and Tena cohorts) originated from stocks established from wild individuals from the same source population in Vilcabamba. The Tena cohort was included, to increase sample size and to further test whether any sex-specific patterns are robust to differences in rearing environment.

Brains were labelled using anti-synapsin immunofluorescence (Ott 2008; Montgomery et al. 2016) imaged with confocal microscopy, and neuropils segmented in Amira-Avizo 2023 (Thermo Fisher Scientific). We measured volumes of key optic and central brain regions previously shown to be ecologically and evolutionarily relevant in *Heliconius* (Montgomery and Merrill 2017; Montgomery et al. 2021), including the medulla, lamina, lobula, lobula plate, accessory medulla, ventral lobula, antennal lobe, anterior optic tubercle, and mushroom body calyx and lobes plus pedunculus. The rest-of-central-brain (rCBR) volume was used as an allometric control. Paired structures were quantified in one hemisphere and doubled, and volumetric data were z-axis corrected using a scaling factor of 1.52 (Montgomery and Ott 2015) and log_10_-transformed before analysis. We measured brain volumes for the 17 newly collected butterflies and combined these data with the previously published measurements for the 26 individuals from Montgomery and Merrill (2017).

### Statistical analysis

All analyses were conducted in R v.4.5.1 (R Core Team 2025). Generalized linear mixed-effects models (GLMMs) and linear mixed-effects models (LMMs) were fitted using the *lme4* package (Bates et al. 2015), and all figures were produced with *ggplot2* (Wickham 2011). Estimated marginal means were computed using the *emmeans* package (Lenth et al. 2019).

Pollen presence was modelled with binomial GLMMs including sex and time of day and their interaction as fixed effects and random effects for individual and date. We used likelihood-ratio tests (LRTs) to assess the significance of sex, time of day and their interaction by comparing each full model to a reduced model lacking the predictor of interest. Pollen load categories were analysed with cumulative link models using the *ordinal* package (Christensen 2015), with sex and time of day as fixed effects. Activity patterns, inferred using time of capture, were analysed by modelling the probability of capturing males versus females as a function of hour of day via multinomial regression (*nnet* package; (Venables and Ripley 2002). Body size was tested with linear mixed models including sex as a fixed effect and observer as a random effects. Movement distances of recaptured individuals were calculated with Vincenty ellipsoid methods implemented in the *geosphere* package (Hijmans 2024), and analysed with log-transformed linear models including sex and body length as fixed effects. Microhabitat associations (Temperature (°C) and relative humidity (%)) with sex were examined with binomial GLMMs including temperature and relative humidity as continuous predictors and date as a random effect.

To test effects of sex, body part (proboscis vs. gut), and predicted age on pollen composition, we used PERMANOVA on Bray–Curtis dissimilarities with the *adonis2* function in vegan (Oksanen et al. 2025). Pollen richness (observed taxa) and diversity (Shannon index) were modelled with GLMMs, including date as a random effect. Age was modelled as a continuous predictor; results were compared with likelihood ratio tests and visualized with estimated marginal means.

Sexual dimorphism in eye and brain morphology was tested using linear models and standardized major axis (SMA) regression. For eyes, log_10_-transformed corneal area and facet number were modelled with sex, log_10_-transformed hind tibia length, and log_10_-transformed interocular width as fixed effects. For neuropils, log_10_-transformed volumes were modelled with sex, rearing location (wild, Cambridge, or Tena), and log_10_ rest-of-central-brain volume (rCBR) as fixed effects. Because our dataset included individuals from three environments but all originating from the same source population, we explicitly tested whether sexual dimorphism was robust to differences in rearing environment by including the Sex × Location interaction in all models. Model residuals were checked for normality, and fixed effects were evaluated with Type II ANOVA. Likelihood ratio tests assessed the contribution of predictors, and post hoc contrasts were performed with *emmeans* using Bonferroni correction. SMA regression (smatr package (Warton et al. 2012)) was used to test sex-specific differences in slopes, intercepts (elevation shifts), and major axis shifts along a common slope. We report test statistics, p-values, and effect sizes as correlation coefficients (r) derived from the reported statistics; for elevation and shift tests, direction indices (DI) are provided where applicable. P-values from SMA elevation and shift tests were adjusted using the Benjamini–Hochberg false discovery rate (FDR) procedure. Results from both linear models and SMA analyses are summarized in Table S1-S2. Mushroom body calyx and lobes plus pedunculus were analysed separately, consistent with previous *Heliconius* studies (Montgomery et al. 2016; Montgomery and Merrill 2017).

## Results

### Sex Differences in Pollen Presence and Load

Our field collections resulted in records from 371 females and 506 males, which we used to test whether the probability of carrying pollen differed between the sexes. Overall, females were more likely to both carry pollen and tend to carry greater quantities compared to males. Likelihood-ratio tests comparing binomial GLMM models with and without each predictor showed that sex had a significant effect on pollen presence (χ² = 16.59, df =1, *P* < 0.001), whereas time of day did not (χ² = 0.46, df = 1, P = 0.50) and the Sex × Capture Time interaction did not improve model fit (χ² = 0.03, df = 1, P = 0.86). Fitting a cumulative link model analysing pollen load categories, “no”, “small” and “large” (Figure 2A), also revealed that females were significantly more likely to carry both small (estimate = 0.09, z = 4.31, P < 0.001) and large pollen loads (estimate = 0.06, z = 4.00, P = 0.0001) than males.

**Figure 2:**
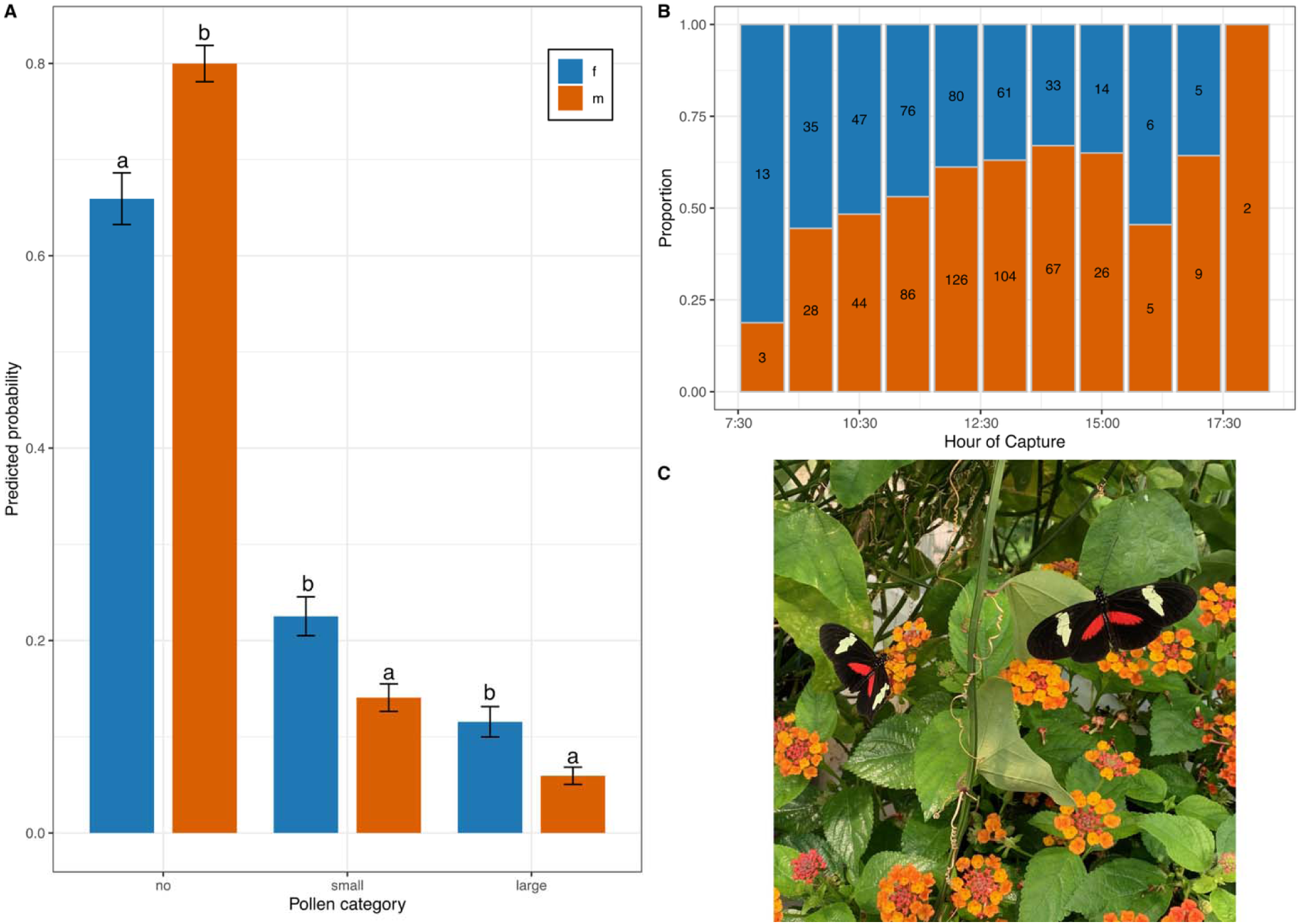
Pollen loads and temporal activity patterns in H. himera. **(A)** Estimated marginal means of predicted probabilities for each pollen load category by sex, derived from a cumulative link mixed model. Letters denote significant pairwise differences between sexes within each category (α = 0.05). (**B** Proportional distribution of female (blue) and male (orange) captures by hour of day, with absolute counts shown inside bar segments. (**C**) H. himera actively foraging on Lantana flowers.

### Pollen richness and diversity differ by body part but not by sex

To investigate the diversity of pollen resources used by *H. himera* and assess potential differences between sexes, sample types, and individual age, we performed DNA metabarcoding on pollen collected from 25 females and 46 males proboscis samples, as well as gut samples from a subset of these individuals (13 females and 26 males). Metabarcoding yielded 1,309,788 high-quality, taxonomically classified reads (mean = 11,907 per sample), detecting 54 families, 122 genera, and 163 species. All samples passed quality filtering and were retained for downstream analysis. The most common taxa included *Lantana camara* (Verbenaceae)(Fig. 2C), *Stellaria media* (Caryophyllaceae), and several Asteraceae species: *Chromolaena odorata*, *Kaunia rufescens*, and *Fleischmannia pycnocephala* (Fig. 3A; Fig S1-S2).

**Figure 3:**
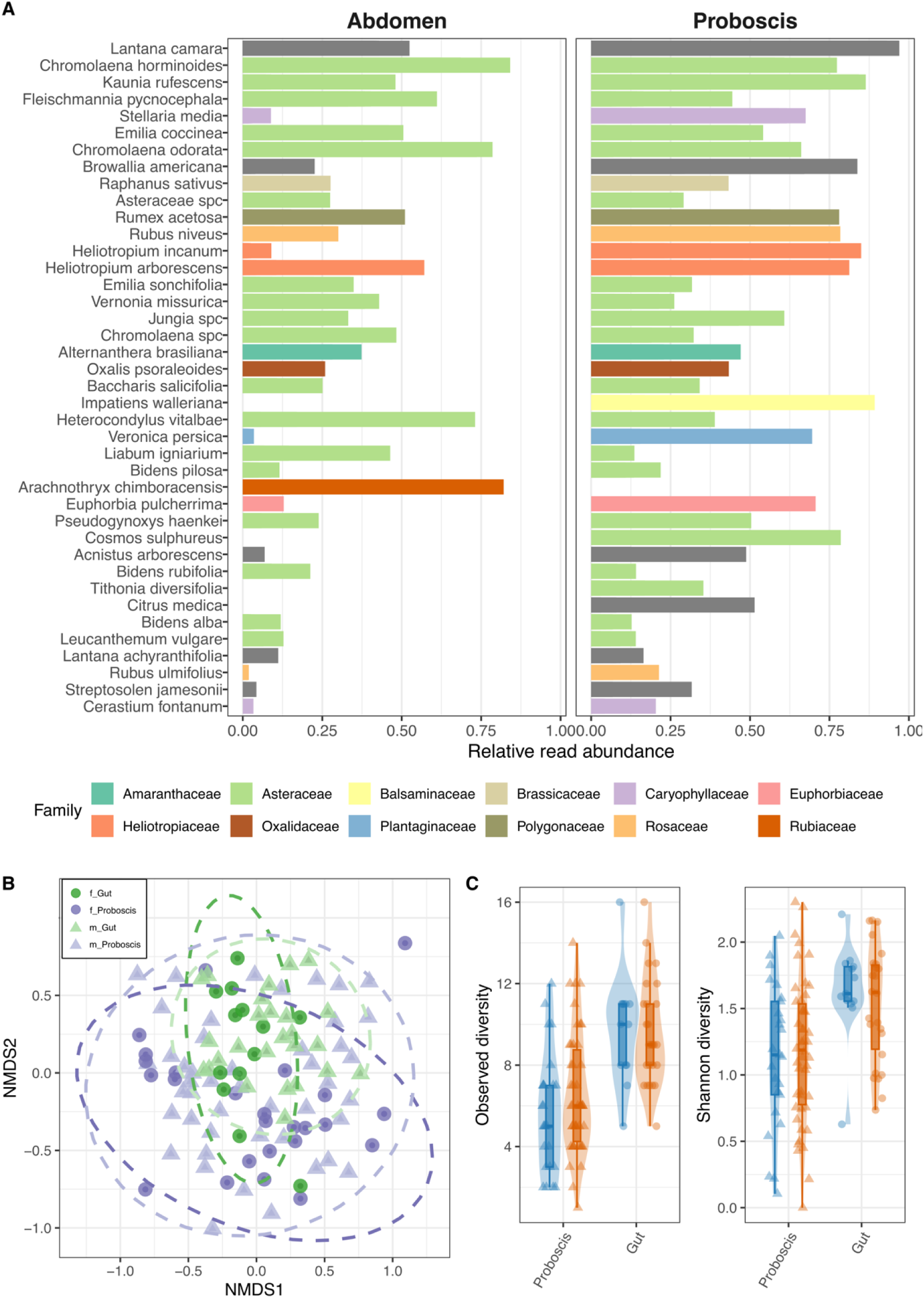
Diversity and composition of pollen collected from different sexes and tissues. (**A**) Relative read abundance of the most common plant taxa identified in proboscis and gut samples. Bars represent the mean relative abundance across individuals, with colors indicating plant families. (**B**) NDMS plot across samples, showing clustering by body part (proboscis = purple, gut = green) and sex (female = circles, male = triangles). 95% confidence ellipses are shown for each group. (**C**) Observed richness and Shannon diversity of pollen samples collected from proboscis and gut tissues of H. himera, separated by sex (females blue and males orange).

Non-metric multidimensional scaling (NMDS) ordination revealed substantial overlap in pollen composition, with no significant differences between sexes (PERMANOVA: R² = 0.0096, F = 1.05, P = 0.378). In contrast, pollen composition differed significantly between body parts (R² = 0.0325, F = 3.63, P = 0.001; Fig. 3B), suggesting some differentiation in the pollen sources detected externally versus internally, possibly reflecting variation in sampling windows or digestion. We found no evidence that predicted age explained variation in pollen composition (R² = 0.014, F = 1.03, P = 0.415; Fig. S3).

Generalized linear and linear mixed models confirmed that observed plant species richness and Shannon diversity differed significantly by body part, with gut samples containing greater richness (χ² = 29.80, df = 1, P < 0.001) and diversity (χ² = 15.56, df = 1, P < 0.001) than proboscis samples (Fig. 3C). In contrast, sex had no effect on either metric (richness: χ² = 1.00, P = 0.32; Shannon: χ² = 0.002, P = 0.97). Predicted age did not influence richness or diversity when modelled as a continuous variable (richness: χ² = 0.19, P = 0.66; Shannon: χ² = 1.78, P = 0.18).

### Daily activity, movement, habitat use and age

Given the sex differences in pollen presence and load size, we next tested for variation in body size, daily activity, movement, and microhabitat use (Fig. S4). Females averaged 24.4 mm in length (SD = 1.66, IQR = 23–26), while males averaged 25.3 mm (SD = 1.55, IQR = 24–26). Linear models confirmed males were larger (χ² = 129.73, df =1, P < 0.001; Fig. S4)

Daily activity also differed between the sexes (LR = 19.11, P < 0.001), with females initiating foraging behaviour earlier in the day (Fig. 2B). To test whether time of day influenced pollen presence, we included capture time as a fixed effect in the binomial GLMM. After accounting for sex (χ² = 16.59, df=1, P < 0.001), capture time did not improve the model (χ² = 0.46, df=1, P = 0.50), and the Sex × Capture Time interaction was also non-significant (χ² = 0.03, P = 0.86). Thus, earlier female activity did not translate into increased pollen loads.

Movement was analysed from 109 recaptured females and 162 males. Recaptured distances varied widely but were comparable: females moved a mean of 87.2 m (median = 24.8, SD = 176.0) and males 66.9 m (median = 36.2, SD = 95.1). A linear model of log-transformed distances showed no effect of sex on movement distance (χ² = 0.63, P = 0.61; Fig. S4).

Microclimate data were available for an opportunistic subset of individuals and were used to explore potential sex differences in local microclimatic conditions. Both sexes were recaptured under similar microclimatic conditions. Females were recaptured at a mean temperature of 25.8 °C (median = 25.9, SD = 3.5) and 50.9% relative humidity (median = 48.0, SD = 16.8), while males were recaptured at 26.4 °C (median = 26.2, SD = 3.4) and 49.5% relative humidity (median = 48.3, SD = 15.0). These distributions overlapped strongly, with interquartile ranges of 23.3–27.9 °C for females and 24.1–29.3 °C for males. Binomial GLMMs confirmed that neither temperature (χ² = 1.05, df = 1, P = 0.31) nor humidity (χ² = 0.11, P = 0.75) significantly affected the probability of capturing either sex (Fig. S4).

### Sexual dimorphism in eye and brain morphology

We tested for sexual dimorphism in eye and brain morphology (Fig. 4). Males had larger eyes than females (F_1,39_ = 26.25, P < 0.001). SMATR analysis showed no sex difference in slope between corneal area and tibia length but a significant elevation shift (Wald χ² = 68.82, FDR-p < 0.001), with males consistently having larger eyes for a given tibia length (Fig. 4A). Facet number was also higher in males (F_1,40_ = 6.07, P = 0.018). Using corneal area as the allometric control, SMATR again detected no sex difference in slope or elevation but a significant major axis shift (Wald χ² = 21.23, FDR-p < 0.001), showing that males more facets (Fig. 4D).

**Figure 4:**
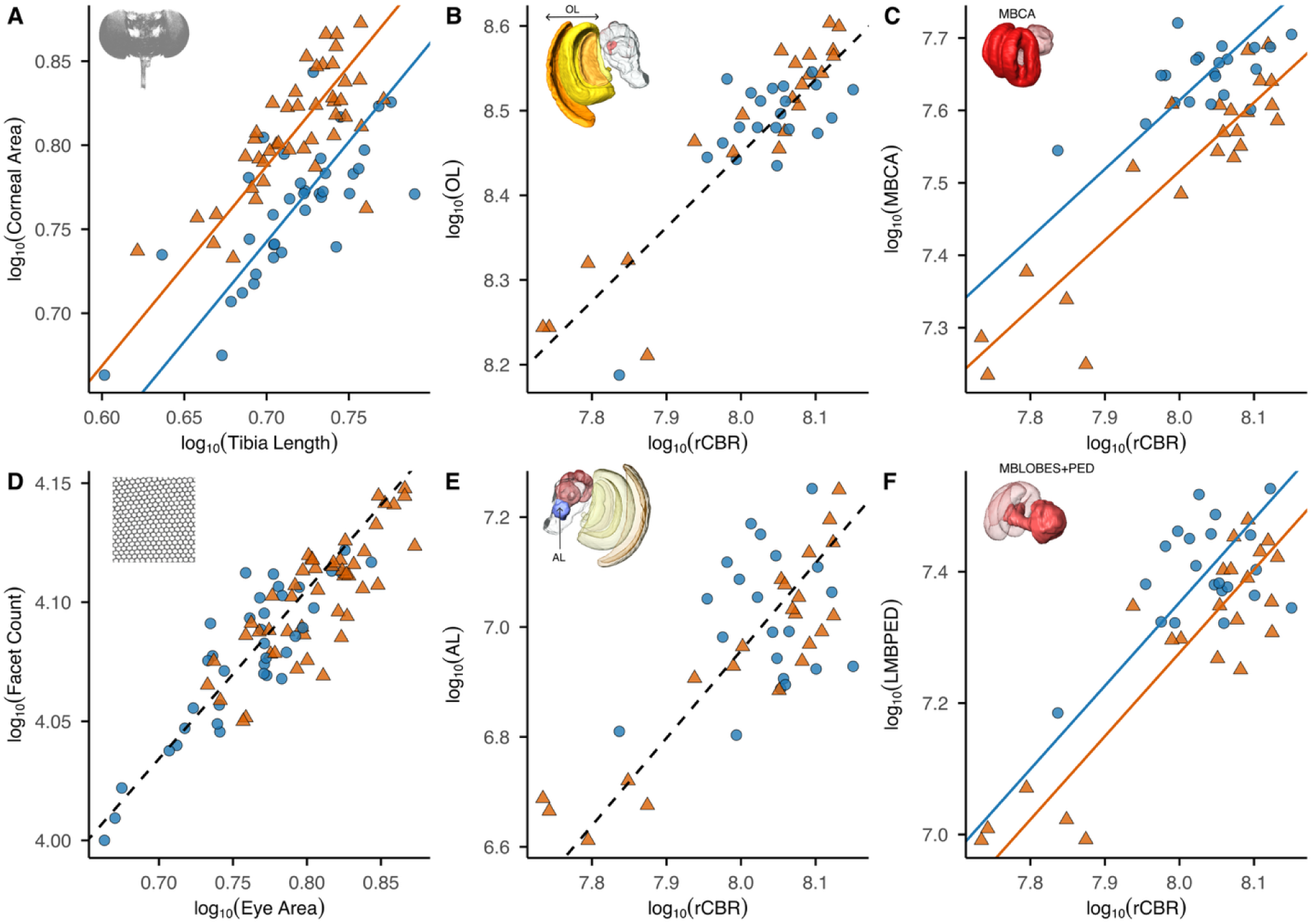
Sexual dimorphism in eye and brain morphology. Females = blue; Males = orange. All traits are log_₁₀-t_ransformed. (**A**) Corneal area shows a significant elevation shift, with males having larger eyes. (**B, E**) Optic lobe (OL) and antennal lobe (AL) volumes scale with rest-of-central-brain volume (rCBR) but show no sex differences. (**C, F**) Mushroom body calyx (MBCA) and mushroom body lobes and pedunculus (MBLOBES+PED) exhibit a significant elevation shift, with females having larger volumes than males. (**D**) Facet count shows a significant major axis shift, with males having more facets. Lines indicate sex-specific SMATR regressions (solid) or common slopes (dashed). Insets show the corresponding anatomical structures: eye (gray), OL (yellow/orange), MB (red), AL (blue).

We also found sex differences in brain morphology, particularly in higher-order integrative regions (Fig. 4B–F). For the mushroom body calyces (MBCA), linear models revealed a strong effect of sex (χ²_₁_ = 28.51, P < 0.001). Rearing location did not significantly affect MBCA volume (χ²_₂_ = 2.81, P = 0.28), and the Sex × Location interaction was also non-significant (χ²_₂_ = 2.99, P = 0.27), indicating that the observed sexual dimorphism was consistent across rearing environments. SMATR analysis confirmed a significant elevation shift in the scaling relationship between MBCA volume and rCBR (Wald χ²_₁_ = 34.25, FDR-p < 0.001), with females having larger MBCA for a given brain size (Fig. 4C). A similar pattern, with larger female investment was observed for the mushroom body lobes and pedunculus (MBPED). Sex showed a significant effect on MBPED volume (χ²_₁_ = 29.38, P < 0.001), with females having larger MBPED. Rearing location showed a main effect (χ²_₂_ = 37.51, P < 0.001), but the Sex × Location interaction remained non-significant (χ²_₂_ = 3.96, P = 0.18). Linear models showed significant effects of sex (χ²_₁_ = 29.5, P < 0.001), after accounting for rCBR and rearing condition location (χ²_₂_ = 49.8, P < 0.0001). SMATR analysis detected a significant elevation shift in the scaling relationship (Wald χ²_₁_ = 8.17, FDR-p =0.023), again indicating larger investment in these mushroom body regions in females (Fig. 4F).

By contrast, we found no evidence of sexual dimorphism in the optic lobes. Sex was not a significant predictor of optic lobe volume (χ²^₁^ = 1.02, P = 0.34), and the Sex × Location interaction was likewise non-significant (χ²_₂_ = 6.19, P = 0.06). Rearing location showed a modest main effect (χ²_₂_ = 8.33, P = 0.02), driven by overall size differences between cohorts, but this did not translate into sex-specific variation (Fig. 4B). Similarly, antennal lobe volumes did not differ between sexes (χ²₁ = 3.92, P = 0.06), and the Sex × Location interaction was not significant (χ²_₂_ = 3.70, P = 0.20). As with the optic lobes, rearing location had a main effect (χ²_₂_ = 52.61, P < 0.001), but this reflected overall size variation rather than sexual dimorphism (Fig. 4E). Full model results are provided in Supplementary Tables S1-S2.

## Discussion

Systematic field sampling combined with DNA metabarcoding revealed high floral diversity and clear sex differences in pollen foraging behaviour in wild *H. himera*. Females carried larger pollen loads than males, consistent with sex-specific nutritional demands; yet we observed no differences we observed no differences plant richness and composition. This indicates that sex differences reflect variation in foraging effort rather than shifts in plant choice. Notably, gut samples contained higher pollen diversity and more consistent community profiles than proboscis samples, suggesting they better capture cumulative foraging history. We further confirmed sexual dimorphism in eye morphology and mushroom body volume, consistent more generally with patterns reported for other aspects of vision across *Heliconius* (McCulloch et al. 2016; McCulloch et al. 2017; Chakraborty et al. 2023; Couto et al. 2023; Wright et al. 2023; Wright et al. 2024; Buerkle et al. 2025; VanKuren et al. 2025). While we did not directly test functional links between sensory traits and foraging effort, our study provides one of the first molecular characterizations of pollen resource use in *Heliconius*, showing that sex-specific foraging reflects differential effort on shared floral resources rather than partitioning of plant species.

The floral resources used by *H. himera* differed markedly from those of lowland *Heliconius*. Metabarcoding revealed a diverse diet of 163 plant species spanning, 122 genera and 54 families. While *Lantana camara* (*Verbenaceae*) was the most frequent resource, *Asteraceae* were also well represented, particularly *Chromolaena* and *Kaunia*, which produce amino acid–rich nectar and are widely visited by butterflies and other pollinators (Wist and Davis 2008; Rathnayake and Wijetunga 2016; Layek et al. 2022). By contrast, *Gurania* and *Psiguria* (*Cucurbitaceae*), central to the foraging ecology of lowland *Heliconius (Estrada and Jiggins 2002; Jiggins 2017)*, are entirely absent in the range and diet of *H. himera*. Instead, *H. himera* relied heavily on *Lantana*, highlighting its role as a core floral resource in montane dry forests. Larval resource use also reflects this environment: *H. himera* oviposits on *Passiflora* species of the *Decaloba* subgenus, which tolerate dry conditions and occur in open habitats (Ulmer and MacDougal 2004). Thus, both larval and adult foraging are shaped by the montane dry-forest environment: larvae exploit drought-tolerant host plants, while adults rely on a broad array of floral resources. Together, these patterns suggest that *erato* clade species, including *H. himera,* have retained a generalist strategy across life stages, facilitating colonisation of high-altitude habitats.

In our study, both sexes of *H. himera* experienced similar environmental conditions and movement patterns, indicating shared microhabitat use and strong spatial overlap in access to floral resources. In patchy, resource-limited montane habitats, such overlap may favour a generalist foraging strategy over sex-specific specialization. Accordingly, sex-specific nutritional demands appear to be met through differences in foraging effort rather than distinct floral species. Females for example, may accumulate larger pollen loads through greater persistence at flowers; recent behavioural work shows that *Heliconius* females often revisit flowers repeatedly, leading to increased pollen accumulation (Dell’Aglio et al. 2024). In other *Heliconius*, microhabitat differences often underlie floral preferences rather than intrinsic species identity (Cardoso 2001; Estrada and Jiggins 2002). For example, *H. melpomene* and *H. cydno* rely on *Lantana* and *Psychotria*, respectively, due to habitat use rather than innate preference (Jiggins 2017). Likewise, long-term studies of *Argynnis idalia* show that sex differences in floral visitation diminish when males and females forage during overlapping phenological windows (Chmielewski et al. 2023). By contrast, *H. charithonia* exhibits clear sex-specific floral preferences (Mendoza-Cuenca and Macías-Ordóñez 2005). The absence of strong floral partitioning in *H. himera* therefore suggests that sex-specific strategies are expressed mainly through pollen collection effort, as an adaptive response to ecological constraints of montane environments.

Our comparison of gut and proboscis samples confirmed that tissue type strongly affects pollen detection. Gut samples showed higher richness and diversity, likely reflecting a longer integration window and cumulative foraging history or more passive uptake from nectar. Because whole abdomens were processed, we cannot exclude the possibility that pollen on the abdomen exterior contributed to gut-sample detections; however, we note that this might still be informative for assessing habitat use. Proboscis samples were often dominated by *Lantana*, suggesting its importance as a core pollen source, actively processed through prolonged proboscis manipulation and digestion. In *Heliconius*, pollen digestion occurs externally on the proboscis surface, and only dissolved nutrients are ingested (Gilbert 1972; Penz and Krenn 2000; Eberhard et al. 2007; Hikl and Krenn 2011; Harpel et al. 2015). As a result, pollen from nectar-only visits or brief contacts may be detected externally, which could explain why some taxa (e.g., *Impatiens walleriana*) appeared exclusively in proboscis samples. Together, these patterns suggest that gut samples may reflect longer-term foraging, whereas proboscis samples more closely reflect recent plant interactions and active pollen processing.

Beyond these tissue-specific insights, our results highlight the power of DNA metabarcoding for ecological research in butterflies. This approach enabled species-level identification of complex pollen mixtures from small tissue samples, detecting a broad taxonomic spectrum not easily resolved by classical palynology (Bell et al. 2016). The successful use of DMSO-preserved samples further demonstrates its potential for archived collections, opening opportunities to compare historical and spatial variation in pollen use, shifts in foraging behaviour, and interspecific differences across the *Heliconius* clade. For example, seasonal or local changes in pollen availability may influence investment in cyanogenic glycosides or egg-laying effort (Jiggins 2017; Young and Montgomery 2020). Leveraging DNA metabarcoding across temporal and environmental gradients can thus deepen our understanding of the ecological and evolutionary dynamics structuring pollinator–plant interactions in tropical ecosystems.

Higher pollen loads in females likely reflect reproductive and physiological demands. This pattern aligns with earlier observations in other *Heliconius* species, where females forage more actively and carry larger pollen loads than males (Boggs et al. 1981; Cardoso 2001; Mendoza-Cuenca and Macías-Ordóñez 2005). Female *Heliconius* require a steady intake of amino acids and lipids for oogenesis and to maintain cyanogenic glycosides that serve as chemical defenses for adults and offspring (Gilbert 1972; Dunlap-Pianka et al. 1977; O’Brien et al. 2003; Pinheiro de Castro et al. 2025). These demands cannot be met through nectar alone, making pollen essential for replenishing protein reserves. Males also benefit from amino acids for spermatophore production and maintenance (Boggs and Gilbert 1979; Boggs et al. 1981), but carried lower pollen loads, possibly reflecting weaker nutritional demands, greater time investment in mate search, or greater efficiency in resource extraction. Female-biased foraging extends beyond *Heliconius*: across Lepidoptera, females preferentially use nutrient-rich resources supplying amino acids, lipids, and sterols required for egg production (Alm et al. 1990; Erhardt and Rusterholz 1998; O’Brien et al. 2004; Mevi[Schütz and Erhardt 2005; Keller et al. 2015; Chmielewski et al. 2023). This sex-specific investment pattern is consistent with optimal foraging theory, where individuals adjust investment in foraging behaviour according to the relative fitness payoffs of nutrient acquisition (Houston 1995). For females, the high returns of pollen amino acids for egg production outweigh the time and energetic costs of prolonged foraging, while males balance pollen collection against investment in mate-searching behaviour. Although females began foraging earlier in the day, time of day did not predict pollen presence, nor did its effect differ between sexes, indicating that earlier activity alone does not explain the observed female bias in pollen load. More broadly, sex-based differences in resource use are an important source of intraspecific variation that may influence pollination networks and community dynamics (Smith et al. 2019; Kishi and Kakutani 2020; Smith et al. 2021).

We used full-spectrum wing reflectance to estimate the relative age of wild butterflies, adapting a model trained on known-age individuals. Although wing wear and colouration are widely used as age proxies in Lepidoptera (Kemp 2006; Dell’Aglio et al. 2017), they are influenced by environmental exposure and behaviour, so reflectance-based estimates should be interpreted cautiously. In our dataset, predicted age did not explain variation in pollen richness or diversity. This contrasts with earlier work on different *Heliconius* species from Trinidad and Costa Rica (Boggs et al. 1981) which reported age-related peaks in pollen load, suggesting that age effects may be weak or context dependent.

Our analyses revealed pronounced sexual dimorphism in the visual system, with males possessing significantly larger eyes and more facets than females. Similar male-biased visual investment has been widely reported in insects and is often associated with mate detection or mate-search behaviour (Ziemba and Rutowski 2000; Somanathan et al. 2017). In contrast, males and females visited the same floral species, indicating that variation in eye morphology does not correspond to differences in floral resource breadth. *H. himera* females had significantly larger mushroom bodies, consistent with the *Heliconius*-wide patterns (Couto et al. 2023), but functional basis of this dimorphism remains unclear: laboratory assays of learning and memory in *Heliconius* have not detected sex effects (Toure et al. 2020; Young et al. 2024; Hodge et al. 2025). We therefore treat the neuroanatomical patterns as descriptive rather than explanatory. Future work should directly test whether males and females differ in colour discrimination, odour sensitivity, or flower-approach behaviour, and whether the observed visual dimorphism is primarily shaped by sexual selection rather than foraging ecology.

Together, our findings show that female *H. himera* carry larger pollen loads than males, yet both sexes exploit a similar diversity and composition of floral resources, indicating sexes adopt divergent behavioural strategies on effort rather than specialization on different floral species. In patchy montane environments, such a generalist strategy may facilitate colonisation of high-altitude habitats. Our use of DNA metabarcoding highlights the value of molecular tools for reconstructing butterfly diets and linking sex-specific foraging to broader questions of intraspecific variation and pollination ecology. Gut samples proved particularly informative, capturing a broader and more integrated record of pollen use than proboscis samples and underscoring the importance of tissue choice in dietary inference. We also documented sexual dimorphism in eye and brain morphology, consistent with patterns across *Heliconius*, although the functional significance of these traits remains unresolved.

## Acknowledgements

We are grateful to the Ministerio del Ambiente, Agua y Transición Ecológica for permission to collect butterflies in Ecuador (MAATE-DBI-CM-2021-0176; MAATE-DNB-CM-2021-0176). We thank Alicia Cerchiai for allowing us access to Bosque y Vegetación Protectores Rumi Wilco. Alicia Cerchiai, Alejandro Mancino, Verónica Romero, and Veronika Schottenheim contributed to plant and butterfly stock maintenance. Yi Peng Toh assisted with brain imaging.

Portions of the text were revised with assistance from OpenAI’s ChatGPT (GPT-4, accessed July 2025) to improve clarity and style. All content was critically reviewed and finalized by the authors.

## Author Contributions

J.B.: conceptualization, data curation, formal analysis, investigation, visualization, writing—original draft, writing—review and editing; D.F.R.S.: conceptualization, investigation, writing—review and editing; D.S.W.: methodology, writing—review and editing; S.H.M.: conceptualization, supervision, writing—review and editing; A. K.: resources, formal analysis, writing—review and editing.; R.M.M.: conceptualization, resources, supervision, writing—original draft, writing—review and editing.

## Data Availability Statement

All datasets, analysis scripts, and metadata associated with this study are publicly available on Zenodo. The full repository can be accessed at: https://doi.org/10.5281/zenodo.17777739

