## Supplementary Material for "Sexual dimorphism in pollen foraging and sensory traits in *Heliconius* butterflies"

Supplementary Material: Sexual dimorphism in pollen foraging and sensory traits in *Heliconius* butterflies
All datasets, analysis scripts, and metadata associated with this study are publicly available on Zenodo. The full repository can be accessed at: <https://doi.org/10.5281/zenodo.17777739>

#### Replication statement:

| **Scale of inference** | **Scale at which the factor is applied** | **Number of replicates** |
| --- | --- | --- |
| Individual butterflies | Sex (female, male)  Eye morphology | 37 females, 48 males |
| Individual butterflies | Sex (female, male)  Brain morphology | 21 females, 22 males |
| Individual butterflies | Sex (female, male)  Pollen load & activity | 371 females, 506 males |
| Individual butterflies | Sex (female, male)  Distance recaptures | 109 females, 162 males |
| Individual butterflies | Sex (female, male) Microclimatic conditions | 24 females, 25 males |
| Individual butterflies | Predicted age group  (via wing reflectance) | 25 females, 46 males |
| Individual butterflies | Sex (female, male)  Pollen composition | 25 females, 46 males |
| Individual butterflies | Body part (gut, proboscis) Paired samples | 71 proboscis (25 females, 46 males); 39 gut (13 females, 26 males) |

### Supplementary Tables

**Table S 1:** Likelihood-ratio tests for effects of Sex, Location, Sex×Location, and rest central brain volume (rCB) on neuropil volumes in Heliconius himera.

| **Neuropil** | **Predictor** | **LogLik_reduced** | **LogLik_full** | **CHISQ** | **p_value** | **Adjusted_p_value** | **Significance** |
| --- | --- | --- | --- | --- | --- | --- | --- |
| AL | Location | 35.35 | 61.654 | 52.608 | 0.000 | 0.000 | *** |
| AL | SexLocation | 63.616 | 65.468 | 3.704 | 0.198 | 0.311 |  |
| AL | rCB | 40.632 | 61.654 | 42.043 | 0.000 | 0.000 | *** |
| AL | sex | 61.654 | 63.616 | 3.925 | 0.057 | 0.089 |  |
| AOTU | Location | 39.211 | 40.213 | 2.005 | 0.394 | 0.394 |  |
| AOTU | SexLocation | 40.949 | 42.274 | 2.649 | 0.319 | 0.350 |  |
| AOTU | rCB | 31.503 | 39.211 | 15.415 | 0.000 | 0.000 | *** |
| AOTU | sex | 39.211 | 39.916 | 1.410 | 0.248 | 0.303 |  |
| LAM | Location | 53.236 | 56.419 | 6.365 | 0.049 | 0.067 |  |
| LAM | SexLocation | 56.419 | 60.876 | 8.916 | 0.016 | 0.059 |  |
| LAM | rCB | 41.818 | 51.757 | 19.878 | 0.000 | 0.000 | *** |
| LAM | sex | 54.641 | 60.876 | 12.472 | 0.007 | 0.019 | * |
| MBPED | Location | 50.986 | 69.741 | 37.509 | 0.000 | 0.000 | *** |
| MBPED | SexLocation | 69.741 | 71.722 | 3.962 | 0.176 | 0.311 |  |
| MBPED | rCB | 48.696 | 71.722 | 46.052 | 0.000 | 0.000 | *** |
| MBPED | sex | 57.033 | 71.722 | 29.378 | 0.000 | 0.000 | *** |
| LOB | Location | 70.543 | 77.158 | 13.229 | 0.001 | 0.002 | ** |
| LOB | SexLocation | 77.158 | 79.626 | 4.936 | 0.112 | 0.246 |  |
| LOB | rCB | 42.642 | 75.333 | 65.382 | 0.000 | 0.000 | *** |
| LOB | sex | 75.333 | 77.158 | 3.650 | 0.067 | 0.091 |  |
| LOP | Location | 66.791 | 74.141 | 14.700 | 0.000 | 0.001 | *** |
| LOP | SexLocation | 74.141 | 80.247 | 12.212 | 0.003 | 0.030 | * |
| LOP | rCB | 53.73 | 80.247 | 53.033 | 0.000 | 0.000 | *** |
| LOP | sex | 73.636 | 80.247 | 13.221 | 0.005 | 0.017 | * |
| MBCA | Location | 66.669 | 68.072 | 2.807 | 0.278 | 0.339 |  |
| MBCA | SexLocation | 68.072 | 69.569 | 2.994 | 0.273 | 0.335 |  |
| MBCA | rCB | 37.883 | 66.669 | 57.572 | 0.000 | 0.000 | *** |
| MBCA | sex | 52.413 | 66.669 | 28.513 | 0.000 | 0.000 | *** |
| ME | Location | 76.762 | 91.508 | 29.492 | 0.000 | 0.000 | *** |
| ME | SexLocation | 91.508 | 95.943 | 8.869 | 0.016 | 0.059 |  |
| ME | rCB | 49.873 | 91.423 | 83.101 | 0.000 | 0.000 | *** |
| ME | sex | 91.423 | 95.943 | 9.039 | 0.038 | 0.070 |  |
| OL | Location | 71.589 | 75.755 | 8.331 | 0.017 | 0.027 | * |
| OL | SexLocation | 75.755 | 78.848 | 6.187 | 0.062 | 0.170 |  |
| OL | rCB | 44.624 | 75.755 | 62.262 | 0.000 | 0.000 | *** |
| OL | sex | 75.247 | 75.755 | 1.015 | 0.341 | 0.375 |  |
| aME | Location | 34.624 | 48.581 | 27.914 | 0.000 | 0.000 | *** |
| aME | SexLocation | 48.863 | 50.354 | 2.983 | 0.274 | 0.335 |  |
| aME | rCB | 38.429 | 48.581 | 20.304 | 0.000 | 0.000 | *** |
| aME | sex | 48.581 | 48.863 | 0.563 | 0.479 | 0.479 |  |
| vLOB | Location | 54.685 | 55.919 | 2.467 | 0.317 | 0.348 |  |
| vLOB | SexLocation | 58.383 | 59.327 | 1.888 | 0.447 | 0.447 |  |
| vLOB | rCB | 35.808 | 57.053 | 42.490 | 0.000 | 0.000 | *** |
| vLOB | sex | 54.685 | 57.053 | 4.736 | 0.031 | 0.068 |  |

**Table S 2**: SMATR tests for sex differences in neuropil allometry in H. himera, including slope shifts (β), elevation shifts (α), and major axis shifts relative to rCB. FDR-corrected p-values, effect sizes (r), and direction of increase (DI) are reported.

|  | **Slope shift (β)** | | |  | **Elevation shift (α)** | | | | |  | **Major axis shift on common axis** | | | | |
| --- | --- | --- | --- | --- | --- | --- | --- | --- | --- | --- | --- | --- | --- | --- | --- |
|  | **Likelihood ratio** | **p** | **FDR** |  | **Wald-statistic** | **p** | **FDR** | **r** | **DI** |  | **Wald-statistic** | **p** | **FDR** | **r** | **DI** |
| ME | 2.268 | 0.132 | 0.295 |  | 0.198 | 0.656 | 0.802 | - | - |  | 0.680 | 0.410 | 0.563 | - | - |
| LAM | 3.816 | 0.051 | 0.186 |  | 3.054 | 0.081 | 0.295 | - | - |  | 0.077 | 0.781 | 0.781 | - | - |
| LOB | 7.991 | 0.005 | 0.052 |  | 1.434 | 0.231 | 0.508 | - | - |  | 0.104 | 0.747 | 0.781 | - | - |
| LOP | 0.039 | 0.843 | 0.903 |  | 0.488 | 0.485 | 0.762 | - | - |  | 1.543 | 0.214 | 0.563 | - | - |
| aME | 1.968 | 0.161 | 0.295 |  | 0.001 | 0.981 | 0.981 | - | - |  | 0.700 | 0.403 | 0.563 | - | - |
| vLOB | 2.173 | 0.140 | 0.295 |  | 2.413 | 0.120 | 0.331 | - | - |  | 2.567 | 0.109 | 0.400 | - | - |
| AL | 1.607 | 0.205 | 0.322 |  | 0.235 | 0.628 | 0.802 | - | - |  | 0.963 | 0.327 | 0.563 | - | - |
| AOTU | 0.102 | 0.750 | 0.903 |  | 0.026 | 0.872 | 0.959 | - | - |  | 1.069 | 0.301 | 0.563 | - | - |
| MBCA | 3.994 | 0.046 | 0.186 |  | 34.253 | 0.000 | 0.000 | 0.986 | female |  | - | - | - | - | - |
| MBPED | 0.015 | 0.903 | 0.903 |  | 8.168 | 0.004 | 0.023 | 0.944 | female |  | - | - | - | - | - |
| OL | 1.028 | 0.311 | 0.427 |  | 1.148 | 0.284 | 0.520 | - | - |  | 0.241 | 0.624 | 0.762 | - | - |

### Supplementary Figures

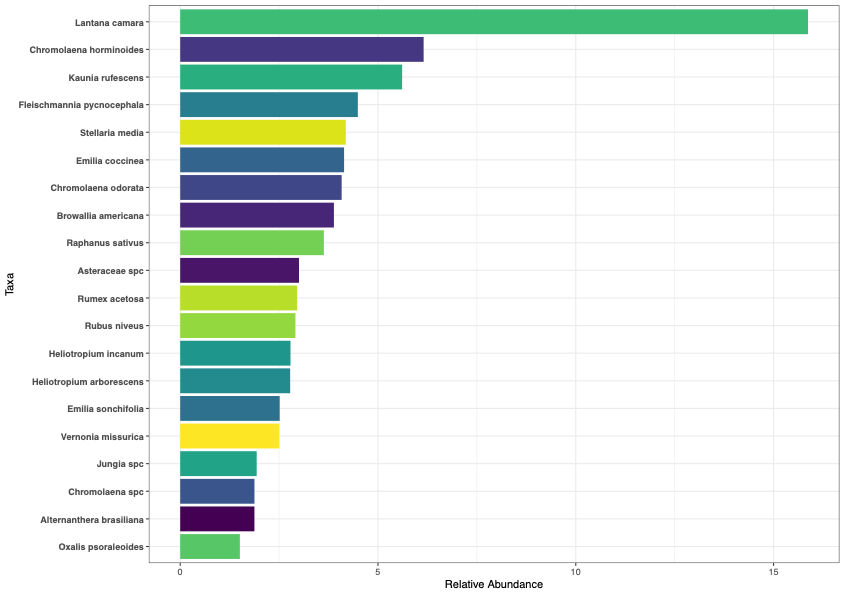

Fig. S 1: **Dominant pollen taxa in H. himera samples.** Barplot showing the mean relative read abundance of the 20 most frequent plant taxa detected across all samples (proboscis and gut combined). Taxa are ordered by abundance.

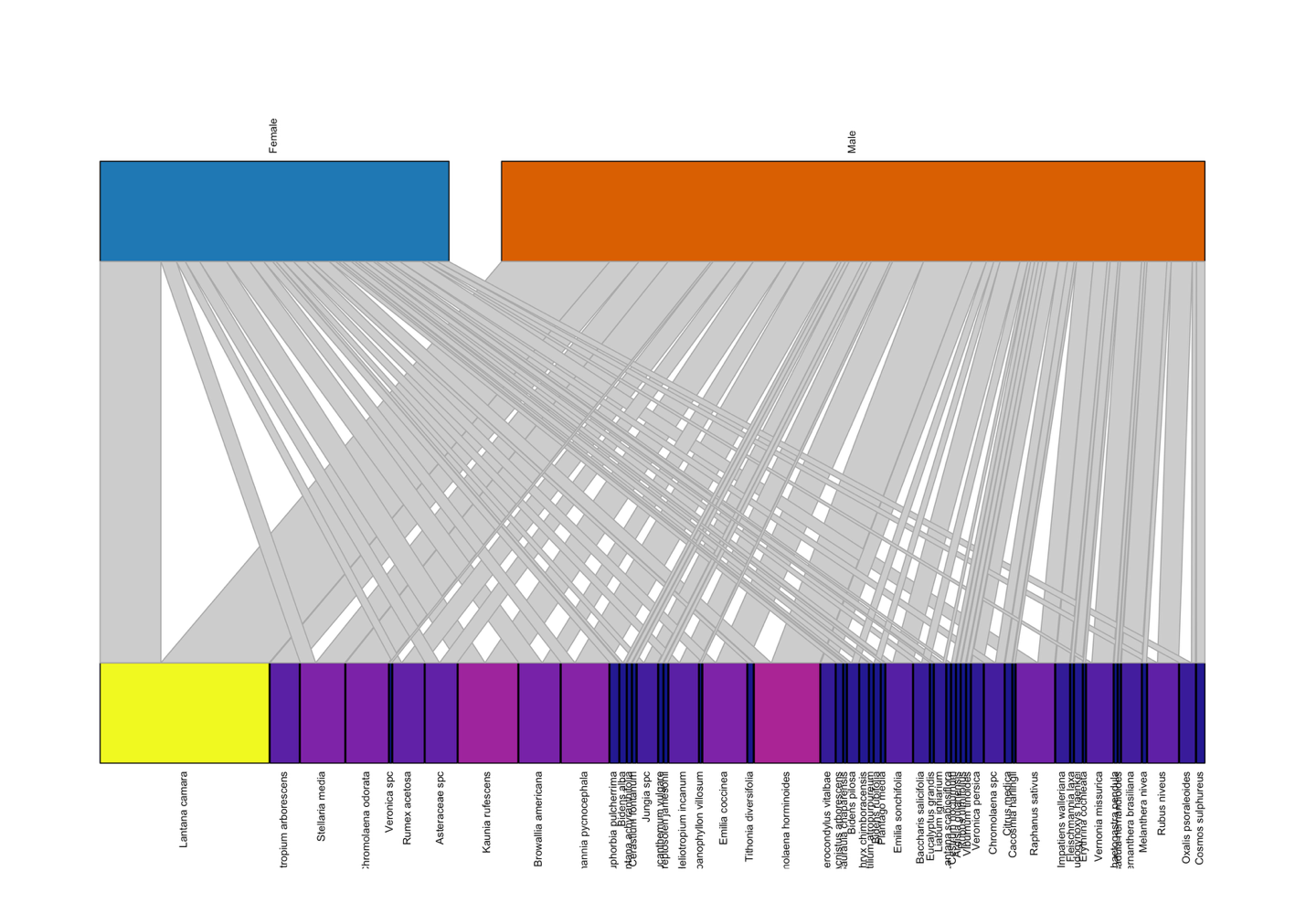

**Fig. S 2: Interaction network between Heliconius himera and the most frequently detected pollen.** Female (Blue) and male (orange) and butterflies are shown at the top, with plant species at the bottom. Edges represent detected interactions, with width proportional to the relative read abundance of each plant in each individual’s pollen load. Edge color indicates interaction strength, ranging from blue (low read count) to yellow (high read count).

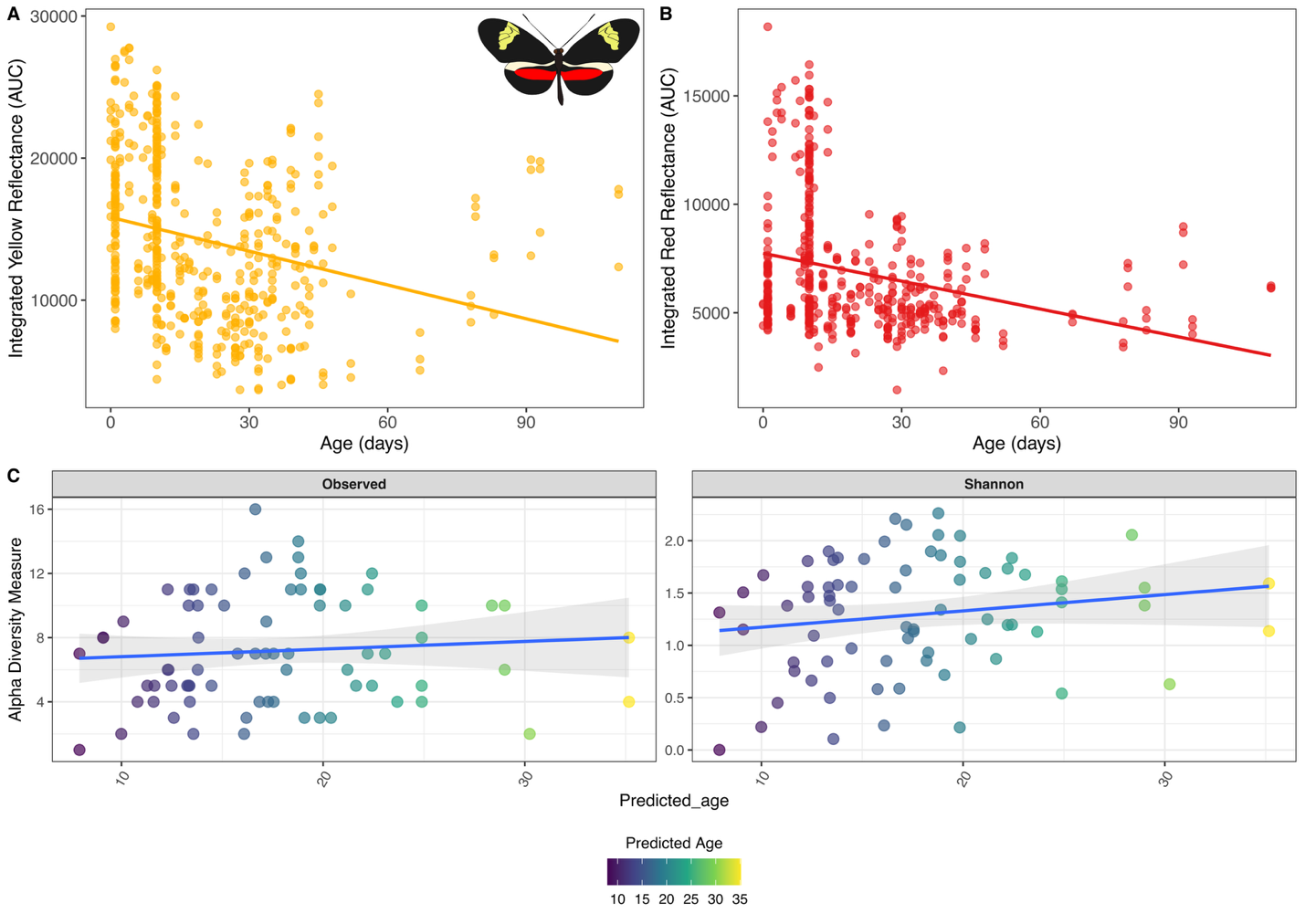

**Fig. S 3: Wing reflectance as an age predictor and its relationship to pollen diversity.** (**A–B**) Spectral reflectance from yellow (**A**) and red (**B**) wing bands of Heliconius butterflies declines with age. Each point represents the integrated area under the reflectance curve (AUC) for an individual butterfly, fitted with a linear model. (**C**) Observed pollen richness and Shannon diversity show a weak positive trend with predicted age based on wing reflectance, but this relationship was not statistically significant. Predicted age was estimated using a model trained on reflectance data from insectary-reared individuals. Color gradient corresponds to predicted age in days.

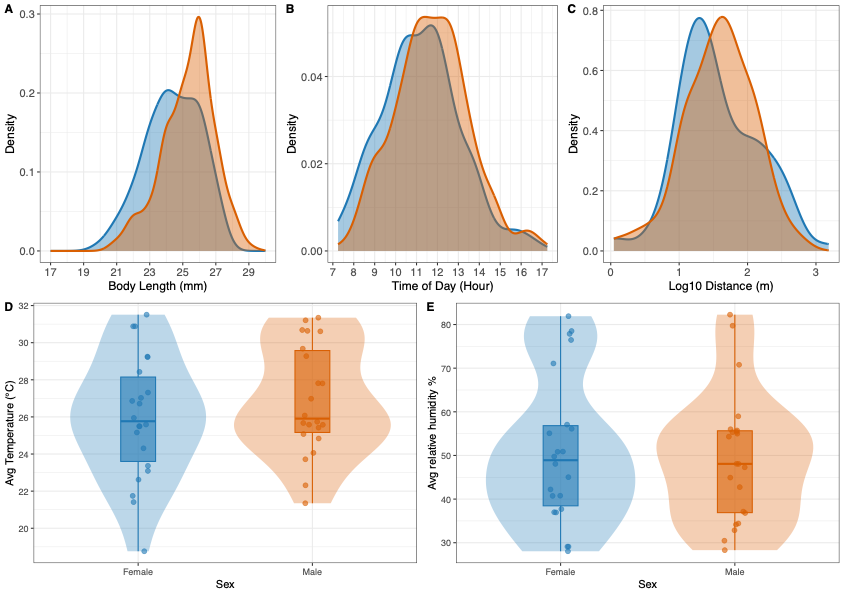

**Fig. S 4: Daily activity, movement and habitat use in Heliconius himera**. (A–C) Density plots showing distributions for body length (A), time of capture (B), and log-transformed distance from capture to recapture site (C) for females (blue) and males (orange). Males tend to be larger and are slightly later during the day, while recapture distances, showed no significant difference between sexes. (D–E) Violin plots with embedded boxplots show average environmental conditions of (**D**) ambient temperature and (**E**) relative humidity. Recapture conditions did not differ between sexes.

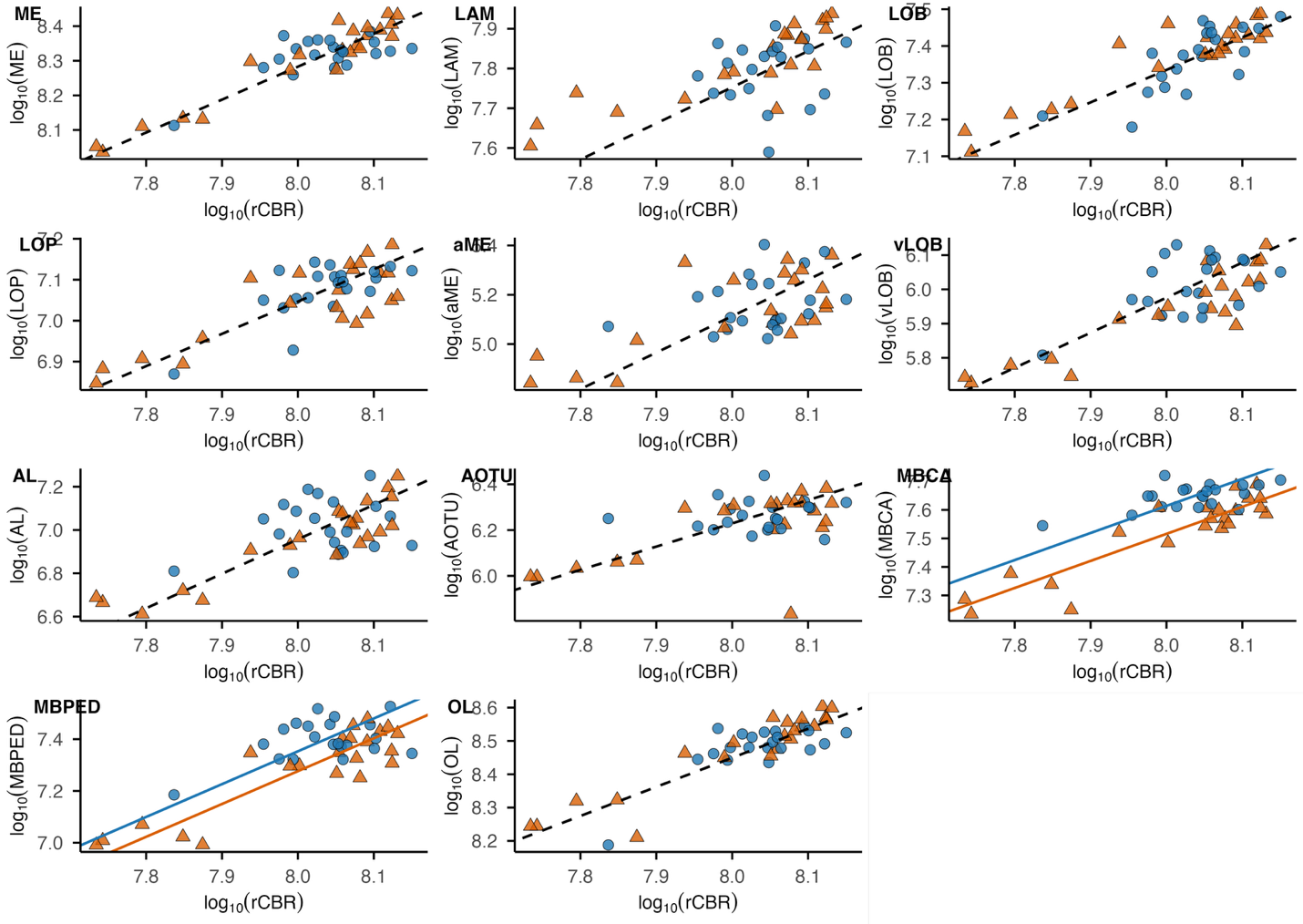

**Fig. S 5: Sexual dimorphism in neuropil scaling relationships in Heliconius himera.** Females (Blue circles) and Males (Orange triangles). All traits are log₁₀-transformed. Pairwise SMATR plots show relationships between individual neuropil volumes and rest-of-central-brain volume (rCB). Lines indicate sex-specific SMATR regressions (solid) or common slopes (dashed).
